# Distinct regulation of collagen biosynthesis by hypoxia and vitamin C uncovers an essential role for CXCL12-CXCR4 signaling

**DOI:** 10.64898/2026.09.14.750894

**Authors:** Méline Ricol, Cindy Dieryckx, Camille Callies, Sandrine Vadon-Le Goff, Jean Baptiste Vincourt, Catherine Moali, Violaine Sée

## Abstract

Collagen biosynthesis is a complex multistep process, essential for functional tissue formation and repair. It is highly active during embryogenesis and wound healing, leading to physiological fibrogenesis, while its dysregulation induces pathologies affecting connective tissues, namely fibrosis. There are several key cellular microenvironmental factors regulating collagen biosynthesis, including vitamin C, oxygen tension and growth factors. Whilst the function of these distinct factors has previously been studied individually, how they operate together, synergistically or not, is not fully characterized. Using primary human dermal fibroblasts, we demonstrated that ascorbic acid (AA) and hypoxia both induced collagen I deposition when applied individually and had additive effects when combined together. Their effects were associated to the expected increase in collagen prolyl hydroxylation. However, AA induced a much stronger collagen secretion compared to hypoxia, while hypoxia leads to a higher collagen I hydroxylation. This observation challenged the common view of collagen prolyl hydroxylation being the central mechanism of AA-induced collagen secretion. Using proteomic analysis of the secretome, biochemical and cell imaging approaches, we further showed that increased CXCL12 extracellular levels and activation of its cognate CXCR4 receptor are at the core of AA-induced collagen intracellular trafficking and secretion. This mechanism is specific to AA and is not involved under hypoxia. The existence of distinct mechanisms underlying the effects of AA and hypoxia is further demonstrated by the different ultrastructure of collagen fibrils in the two conditions. Together, our results contribute to explain how these two triggers drive collagen biosynthesis and pave the way for potential pro- or anti-fibrotic strategies aimed at improving tissue repair.

## Introduction

Fibrillar collagens are abundant and versatile proteins controlling the 3D organization and mechanical properties of tissues and providing important cellular cues. Among these collagens, collagen I is the predominant form and a major component of connective tissues. It is made of a long central triple helical (collagenous) domain characterized by the repetition of the Gly-Xaa-Yaa amino acid sequence, where Xaa and Yaa are mostly proline or 4-hydroxyproline residues respectively [1–3]. The collagenous domain is flanked by two connecting sequences, called N- and C-telopeptides, and two propeptides (one on each extremity) [1–3]. The amount and organization of mature collagen I, deposited and incorporated into the extracellular matrix (ECM) after the proteolytic removal of propeptides, govern tissue stiffness, tensile strength, and architecture. Collagen I synthesis and deposition is therefore central to the structure, mechanics, and function of the extracellular matrix (ECM) in tissues. Moreover, deregulated and excessive accumulation of collagen I is a hallmark of fibrosis in multiple organs, including the lung, liver, kidney, skin and heart. Fibrosis leads to tissue stiffening, disruption of normal organ architecture, and progressive loss of function. Hence, understanding the mechanisms that regulate collagen I deposition is essential for identifying therapeutic targets in fibrotic diseases. In this study, we focused on dermal fibroblasts, which are the main collagen producers in skin. Controlled collagen deposition is key to skin homeostasis and to healthy wound healing. Indeed, insufficient synthesis impairs collagen renewal, tissue strength and wound closure after injury, whereas excessive or aberrant deposition may result in skin thickening and fibrotic scar formation.

Collagen I biosynthesis is a complex and multistep process involving numerous actors localized both in the intracellular and extracellular compartments. Upon transcription and translation, the procollagen α-chains – precursors of collagen – experience multiple essential post-translational modifications within the endoplasmic reticulum (ER), including the frequent 4-hydroxylation of prolines in the collagenous domain. This step is critical to ensure thermal stability, proper folding, fibril formation, and to facilitate collagen secretion and interactions with other ECM components [4–7]. The reaction is catalyzed by the collagen proline 4-hydroxylases (C-P4H1-3) [4–7]. Other important modifications such as proline 3-hydroxylations, lysine hydroxylations and glycosylations also occur within the ER [8–11]. Subsequently, the α-chains assemble into large trimers that are packaged into enlarged vesicles to exit the ER and traffic through the Golgi apparatus before being secreted into the extracellular space [12]. The soluble and immature procollagens require an additional proteolytic maturation, which consists in the release of the N- and C-terminal propeptides and occurs in the late secretory pathway or extracellularly [13]. Together, these steps lead to the integration of mature collagen molecules into the extracellular matrix, where the subsequent formation of cross-links modulates the stiffness of the ECM [8,14]. This final step is controlled by lysyl oxidases that initiate the process by transforming the amine groups of specific lysines into reactive aldehydes.

The C-P4H enzymes, catalyzing proline 4-hydroxylation, are 2-oxoglutarate-dependent dioxygenases (2-OGDD) requiring for their activity, 2-oxoglutarate and oxygen as co-substrates and Fe^2+^ and ascorbic acid (AA) as co-factors [5,15]. These co-substrates and co-factors are also needed by other enzymes involved in collagen biosynthesis, such as collagen proline 3-hydroxylases and lysyl hydroxylases, highlighting their importance in this pathway. These post-translational modifications are essential for collagen stability and incorporation into the ECM, and must therefore be tightly controlled. It places the prolyl hydroxylase coactivators, namely AA and oxygen, at the forefront of a healthy and well-balanced collagen deposition (for review [16]). AA, commonly known as Vitamin C, is a vital organic compound that cannot be synthetized by humans, and thus, must be ingested to sustain the associated metabolic reactions, including collagen synthesis [17]. Its critical role is exemplified by the development of scurvy, in case of AA chronic deficiency, with symptoms associated to dysfunctional collagen [18]. AA is also commonly used in *in vitro* studies related to collagen production and matrix deposition to ensure adequate collagen fibrogenesis. While its function as an enhancer of collagen hydroxylation is well described [19–22], the precise mechanisms by which AA promotes collagen secretion and deposition remain incompletely characterized.

Interestingly, AA also acts on the prolyl hydroxylases that modify the hypoxia inducible factor (HIF) and are key regulators of downstream signaling. Depending on HIF hydroxylation state by HIF-P4Hs, it is either degraded by the proteasome or stabilized and active to induce the transcription of its target genes. Among the well-known HIF target genes are the C-P4Hs [23–25], directly linking the hypoxia signaling pathway to the promotion of collagen deposition [26,27]. As the effects of hypoxia on ECM deposition and remodeling drive the progression of several pathological conditions [28–33], especially cancer and fibrosis, understanding how AA and hypoxia contribute synergistically or not to collagen biosynthesis is relevant for the development of therapeutic tools to fight fibrotic diseases as well as tumor progression.

In this study, we compared the effects of AA and hypoxia, used independently or together, on the collagen biosynthetic pathway activated in skin fibroblasts. We confirmed their common ability to promote collagen proline hydroxylation and demonstrated modest synergistic effects of AA and hypoxia on collagen deposition. Importantly, we also highlighted the different mechanisms by which they promote collagen secretion. Using proteomic analysis, we uncovered an unknown, yet necessary role of AA in promoting the increase of CXCL12 secretion and the activation of its cognate CXCR4 receptor. This specific role of AA compared to hypoxia is associated with enhanced intracellular collagen trafficking and distinct ultrastructure of collagen fibrils. Together, this study describes a new mechanism for AA, in addition to the well-described increased collagen hydroxylation, which is key to efficient collagen secretion and integration into the ECM.

## Results

To elucidate how AA and hypoxia affect collagen I deposition, independently or in conjunction, we used human dermal fibroblasts (HDFs) from five adult donors. We analyzed the impact of the two microenvironmental factors on the intracellular and extracellular maturation mechanisms and on the resulting levels of collagen I deposited into the ECM. Typical culture conditions used 2% Fetal Calf Serum (FCS) to maintain a healthy HDF phenotype, sustain efficient collagen synthesis, and facilitate the analysis of cell supernatants by Western blot, by minimizing serum components in supernatant samples, compared to the classical 10% FCS cultures. Similarly, for hypoxic conditions, 1% O_2_ was used, to trigger an efficient hypoxia signaling, without affecting cell survival or proliferation.

### AA and hypoxia are each sufficient to trigger collagen biosynthesis independently, and also act synergistically

To study the impact of AA and hypoxia on collagen I deposition, HDFs were either cultured in normoxia (21% O_2_) or in hypoxia (1% O_2_), in the presence or absence of a stabilized form of AA (2-phospho-L-ascorbic acid). We first validated the hypoxic response by assessing the expected stabilization of HIF-1α and HIF-2α (Supplemental Figure 1A and 1B) and the increased transcription of representative HIF-target genes upon 24h of hypoxic exposure (Supplemental Figure 1C). Regarding collagen I deposition, unsurprisingly, AA treatment led to increased collagen deposition into the ECM, within 4 days of culture (Figure 1A). Interestingly, hypoxia also increased collagen levels, even in the absence of AA. Collagen I immunostaining showed that AA enhanced collagen I deposition by 3- to 9-fold, depending on the donor, while hypoxia increased collagen I deposition by 2- to 5-fold. When the two stimuli were added simultaneously, collagen deposition was further increased, reaching a 5 to 13-fold increase compared to the normoxic control condition, thereby showing additive effects (Figure 1A). Western blot analysis of the ECM compartment (after decellularization) confirmed the increased levels of mature collagen, by approximately 5-fold, in the presence of each individual trigger and by 8-fold when added together (Figure 1B). This additive effect is surprising, as AA is expected to enhance HIF proteasomal degradation through increased hydroxylation [34–37], and one would rather anticipate an inhibitory effect of AA on hypoxia-induced collagen deposition. Yet, HIF levels and HIF target gene expression were unaltered upon AA treatment in our culture conditions (Supplemental Figure 1).

**Figure 1.**
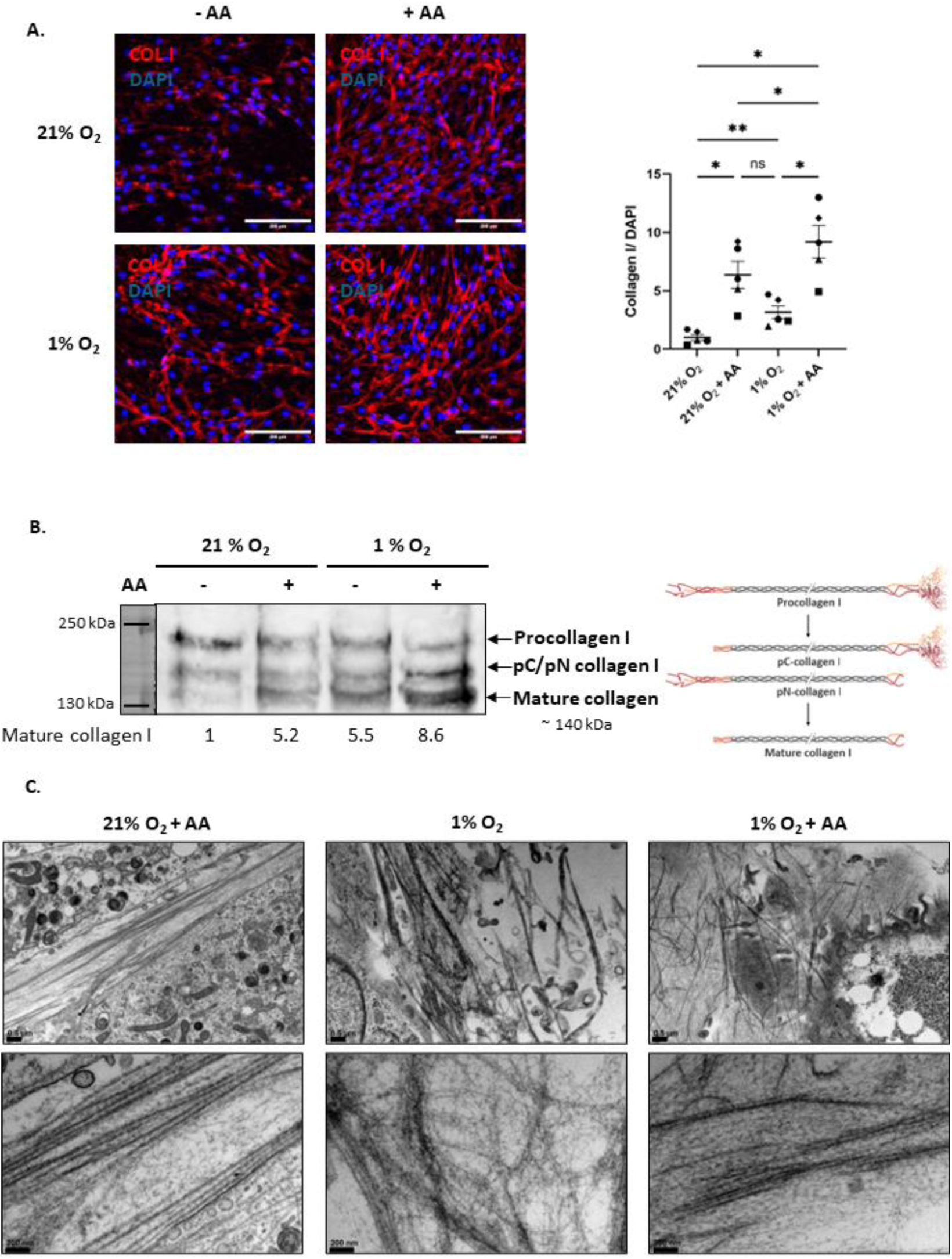
Both AA and hypoxia induce collagen deposition, yet leading to different ultrastructure. Primary HDFs were cultured in normoxia (21% O_2_) or hypoxia (1% O_2_), in presence or absence of stabilized AA (300 µM), as indicated. **A.** Immunofluorescent staining and epifluorescence imaging of collagen I on unpermeabilised fixed fibroblasts after 4 days of culture (red) and DAPI nuclear staining (blue). Quantification was performed with ImageJ; the area covered by collagen I staining was normalized to the area covered by DAPI staining. The results were normalized to the mean of the control condition (21% O_2_). Statistical analysis was performed on N = 5 donors, using RM one-way ANOVA test (normal distribution of data) (ns: non-significant, * p-value < 0.05, ** p-value < 0.01). Scale bar corresponds to 200 µm **B.** Western blot analysis (reducing conditions; 8% acrylamide gel) of collagen I immature and mature forms (depicted on the right) in decellularized ECM protein extracts, after 4 days of culture. Quantification was performed with ImageQuantTL and normalized to the control condition (21% O_2_). **C.** Collagen ultrastructure visualized by TEM after 11 days of culture. Scale bars correspond to 0.5 µm for the upper images and 200 nm for the lower images.

When observed by transmission electronic microscopy (TEM), the ultrastructure of the fibrils looked different depending on the treatment. When treated with AA, the cells produced thick, elongated and well-organized collagen fibres with apparent periodic patterns. However, when cultured under hypoxia (without AA), the fibrils were thinner, shorter and looked less organized. When both AA and hypoxia were applied, the fibrils had a similar appearance to AA only (Figure 1C and Supplemental Figure 2). Whilst these observations are only qualitative, these images suggest that distinct molecular mechanisms might be activated by the two triggers, leading to different ECM organization, and that these mechanisms could be additive.

### Increased collagen I deposition is due to increased collagen secretion

To elucidate the molecular mechanisms involved, we first checked a potential transcriptional activation by AA and / or hypoxia. However, despite the fact that previous studies had demonstrated an increase in collagen I transcription upon hypoxic culture [38–42], neither AA nor hypoxia had a significant impact on *COL1A1* or *COL1A2* transcription in our culture conditions (Figure 2A). AA increased collagen transcription by ∼2-fold in some donors but it varied between donors and did not reach statistical significance. To measure the corresponding protein levels and obtain an unbiased overview of global changes in the entirety of ECM proteins (also known as “matrisome” [43]), we performed mass spectrometry analysis of secreted proteins in the culture supernatant. After 4 days of culture, we found that 23 matrisome proteins were upregulated and 30 were downregulated by AA. The deregulated proteins belonged to all matrisome subcategories, i.e. collagens, proteoglycans, glycoproteins, secreted factors, ECM regulators and ECM-affiliated proteins. Hypoxia led to the deregulation of a similar number of matrisome proteins with 21 upregulated and 36 downregulated proteins (Figure 2B).

**Figure 2.**
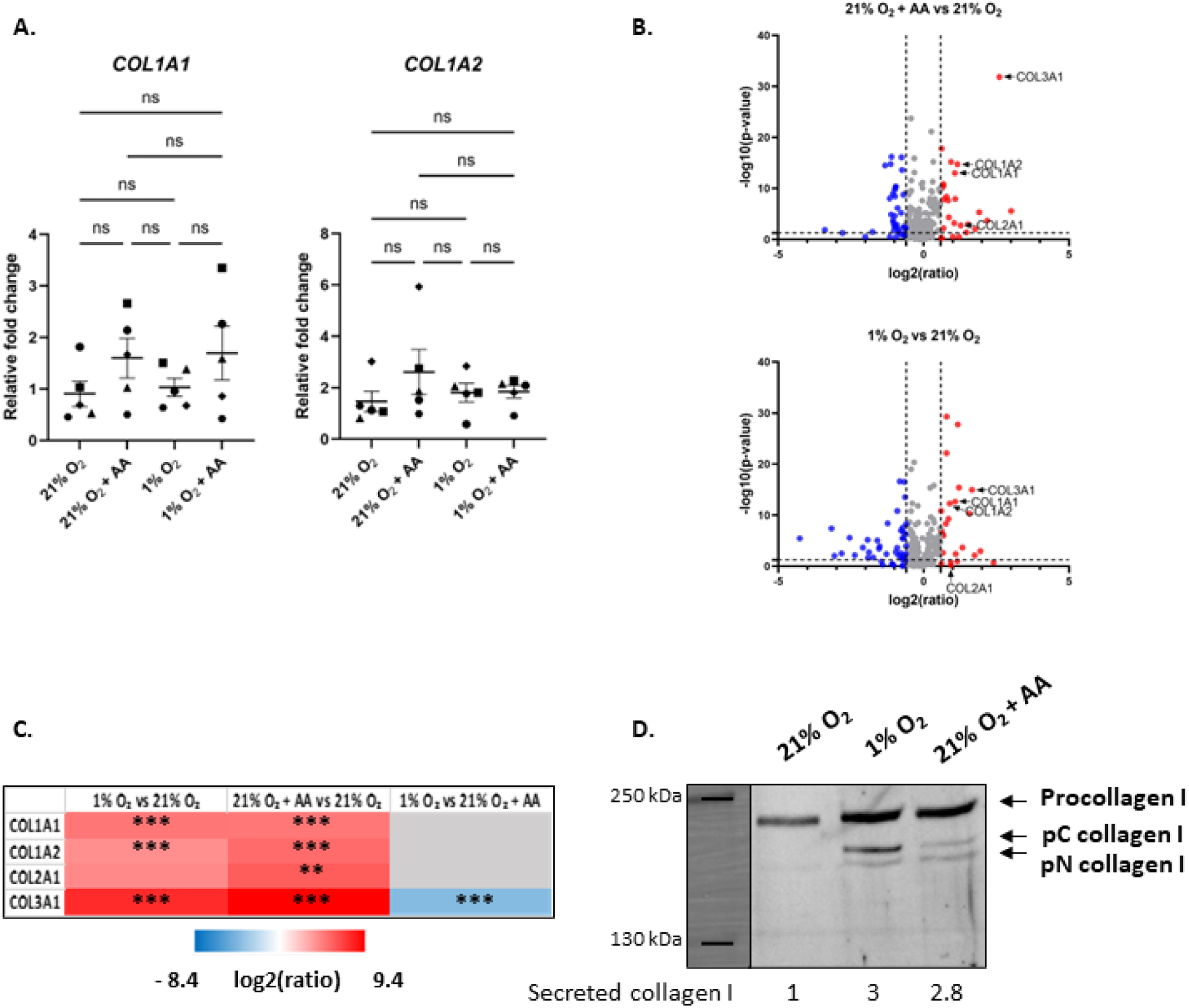
AA and hypoxia induce collagen secretion without increasing its transcription. Primary HDFs were cultured in normoxia (21% O_2_) or hypoxia (1% O_2_), in presence or absence of stabilized AA (300 µM), as indicated. **A.** RT-qPCR analysis of *COL1A1* and *COL1A2* transcripts after 4 days of culture. The results were normalized to the mean of the control conditions (21% O_2_). Statistical analysis was performed on N = 5 donors, using Friedman test (non-normal distribution of data) or RM one-way ANOVA test (normal distribution of data) (ns: non-significant). **B, C.** Supernatants of dermal fibroblasts cultured for 4 days were analyzed using data-independent acquisition mass spectrometry. **B.** Volcano Plots showing up- and down-regulated proteins in indicated conditions (cut-off were set at 1.5 or 0.66 for the fold change and at 0.05 for the p-value). **C.** Heat map of the mass spectrometry data analysis of major fibrillar collagen chains. Protein quantification was carried out on N = 3 donors using a label-free quantitation approach based on fragment ion intensities. The proteins were considered as differentially expressed between 2 conditions when the fold change was > 1.5 or < 0.66, with a p-value < 0.05 (* p-value < 0.05, ** p-value < 0.01, *** p-value < 0.001). **D.** Western blot analysis of cell culture supernatants, collected after 4 days of culture (reducing conditions; 5% acrylamide gel). Quantification was performed with ImageQuantTL and normalized to the control condition (21% O_2_).

We first focused on secreted major fibrillar collagen molecules, and found increased levels (by at least 2-fold) of collagen chains: I α1, I α2 and III α1, in the supernatants of cells treated with AA or hypoxia (Figure 2C). AA treatment also increased the secretion of collagen II α1 while the similar trend in hypoxia did not reach statistical significance (Figure 2B and 2C). Collagen III α1 levels were lower in the supernatants of hypoxic cultures compared to AA-treated cultures, yet this was not the case for collagen I α1 and α2, whose extracellular levels were similar between both conditions (Figure 2C). Finally, when cells were exposed to both hypoxia and AA treatments, secreted collagen levels were similar to the AA only-treated condition (Supplemental Figure 3). Western blot detection of collagen I α1 (Figure 2D) confirmed the mass spectrometry data and showed similar levels of collagen I in the supernatant of cells cultured under hypoxia or treated with AA.

To relate extracellular secreted collagen levels to its intracellular production, we sought to measure intracellular procollagen levels. While the collagen I levels in the supernatants were similar, regardless the pro-fibrogenic culture conditions (AA or hypoxia), the intracellular collagen I contents were strikingly different. Indeed, in the presence of AA, the levels of intracellular procollagen I were much lower compared to hypoxic cultures or to non-treated control conditions (Figure 3A). This suggests that AA promotes a fast and efficient collagen I secretion, which does not occur to the same extent in hypoxic conditions without AA. This strong effect of AA is correlated with a change in the morphology of the rough endoplasmic reticulum (RER). In the absence of AA, we observed numerous large, dark and dilated cisternae, which are likely associated with intraluminal accumulation of proteins within the RER (Figure 3B). On the contrary, AA-treated cells displayed fewer and thinner RER cisternae. This was also the case in hypoxic cell cultures treated with AA, and highlights a key and novel role for AA in maintaining RER homeostasis.

**Figure 3.**
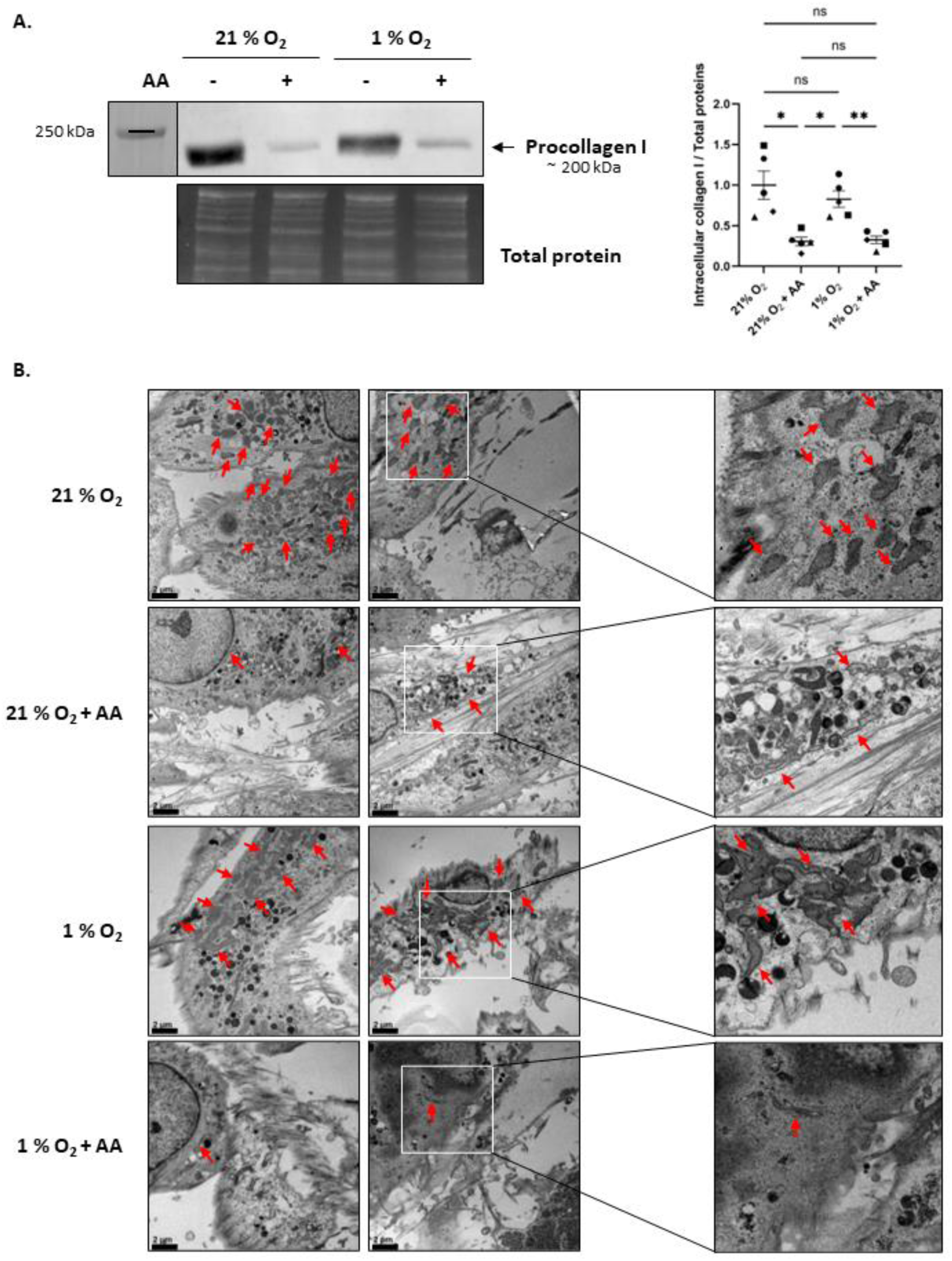
AA induces a faster collagen secretion than hypoxia, reflected by low intracellular levels of procollagen and a differing morphology of the RER. Primary HDFs were cultured in normoxia (21% O_2_) or hypoxia (1% O_2_) in presence or absence of stabilized AA (300 µM) as indicated **A.** Western blot analysis of the intracellular lysates (reducing conditions; 5% acrylamide gel). Procollagen I levels were normalized to total protein levels (stain-free detection) and to the mean of the control conditions (21% O_2_). Statistical analysis was performed on N = 5 donors, using RM one-way ANOVA test (normal distribution of data) (ns: non-significant, * p-value < 0.05, ** p-value < 0.01). **B.** RER ultrastructure visualized by TEM. Scale bar indicates 2 µm.

### AA-induced collagen I secretion cannot be solely explained by increased collagen proline 4-hydroxylation

We hypothesized that the stronger effects of AA on collagen I secretion compared to hypoxia, could be explained by a different level of collagen proline 4-hydroxylation. As mentioned above, collagen proline 4-hydroxylation is a critical step allowing triple helix stabilization and proper folding at body temperature, and facilitating their secretion [6,7]. Both AA and hypoxia are known to increase collagen proline 4-hydroxylation, yet by different mechanisms. AA, as a cofactor of C-P4Hs, maintains reduced iron available for the enzymatic reaction and prevents the inactivation of the enzyme, therefore markedly increasing the rate of collagen proline 4-hydroxylation [19–22]. On the other hand, *P4HA1* and *P4HA2*, the genes encoding C-P4H1 and 2, have been shown to be HIF target genes [24,25,44–46], meaning that hypoxia increases the transcription of these enzymes (for review [16]). We here confirmed that the transcription of *P4HA1*, the most ubiquitously expressed isoform [5], was enhanced by 3-fold in hypoxic conditions (Figure 4A). To compare the efficiency of these two mechanisms on collagen proline 4-hydroxylation (increased enzymatic activity versus increased enzyme levels), we determined the degree of hydroxylation of the collagen molecules in the different conditions. As expected, both AA and hypoxia significantly increased collagen proline 4-hydroxylation, yet with a significantly higher rate for hypoxic culture compared to AA-treated cells. In addition, when both were added together, the hydroxylation levels were even higher (Figure 4B), consistent with the additive effects observed on collagen deposition (Figure 1A). The main conclusion from this experiment is that the different effect of AA and hypoxia on proline 4-hydroxylation cannot be the only mechanism triggering AA-induced collagen secretion. As a matter of fact, the effect of AA is stronger than that of hypoxia on secretion but weaker when looking at collagen proline 4-hydroxylation levels. This suggests that one or several additional mechanisms are required to explain the strong impact of AA in promoting collagen secretion.

**Figure 4.**
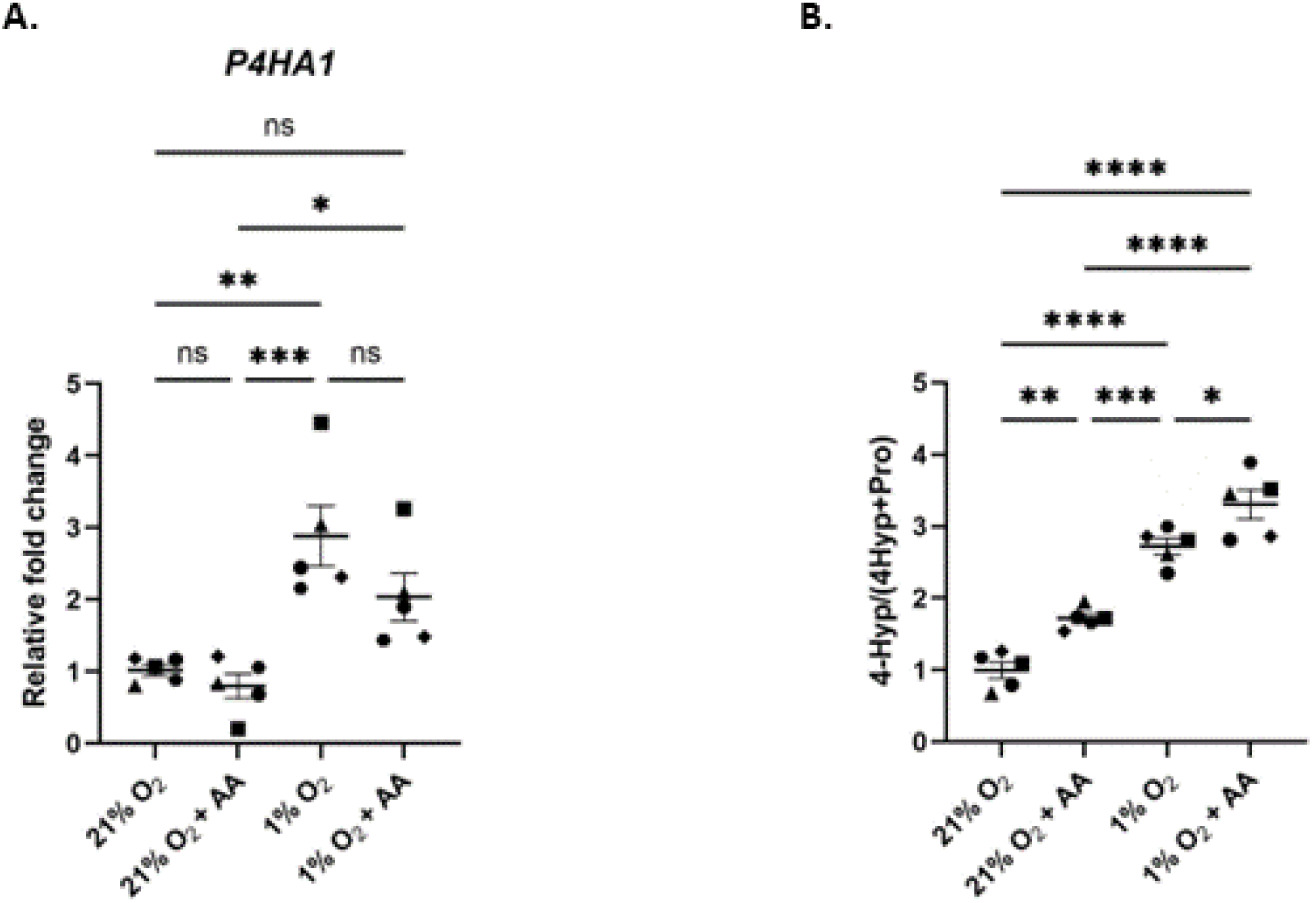
Both AA-and hypoxia induce collagen proline 4-hydroxylation. HDFs were cultured for 4 days in normoxia (21% O_2_) or hypoxia (1% O_2_), in presence or absence of stabilized AA (300 µM), as indicated **A.** RT-qPCR analysis of *P4HA1* transcripts. The results were normalized to the mean of the control condition (21% O_2_). Statistical analysis was performed on N = 5 donors, using RM one-way ANOVA test (normal distribution of data) (ns: non-significant, * p-value < 0.05** p-value < 0.01, *** p-value < 0.005). **B.** Amino acid analysis by MS-UPLC of the hydroxylation rate of extracellular proteins in supernatants. The results were normalized to the mean of the control condition (21% O_2_). Statistical analysis was performed on N = 5 donors, using RM one-way ANOVA test (normal distribution of data) (* p-value < 0.05, ** p-value < 0.01, *** p-value < 0.005, **** p-value < 0.001).

### CXCL12 secreted levels are increased in AA-treated cells but not in hypoxic cells

To depict the additional mechanisms beyond collagen hydroxylation, we went back to mass spectrometry data, and searched for differentially secreted proteins between the 2 treatments. We found 31 proteins belonging to the matrisome, which had changes in abundances in AA-treated cells compared to control non-treated normoxic cells, but which were not affected in hypoxic cultures in comparison to the same control cells (Figure 5A). Among these 31 AA-specific proteins, we identified a subset of 13 proteins, whose abondance also differed significantly after AA treatment of hypoxic conditions (Figure 5B). Among these 13 proteins, we searched for those which had previously been described to be involved in collagen regulation. One promising candidate was CXCL12, which had been shown to play a role in collagen secretion through the enhancement of the formation of large COPII vesicles [47]. CXCL12 secreted levels were 1.6-fold higher in the supernatants of AA-treated cells in comparison to normoxic or hypoxic conditions without AA (Figure 5B). Using ELISA, we confirmed that secreted CXCL12 levels were significantly higher after AA treatment, in comparison to normoxic or hypoxic cultures without AA (respectively 1.7- and 3-fold increase) (Figure 5C). As CXCL12 is a chemokine, that is commonly induced at transcriptional levels, we checked if AA-could up-regulate CXCL12 transcription. Surprisingly, *CXCL12* transcription was not affected by AA (Figure 5D), suggesting that the increased levels of CXCL12 in the supernatant of AA-treated cells were not due to a transcriptional activation of the chemokine.

**Figure 5.**
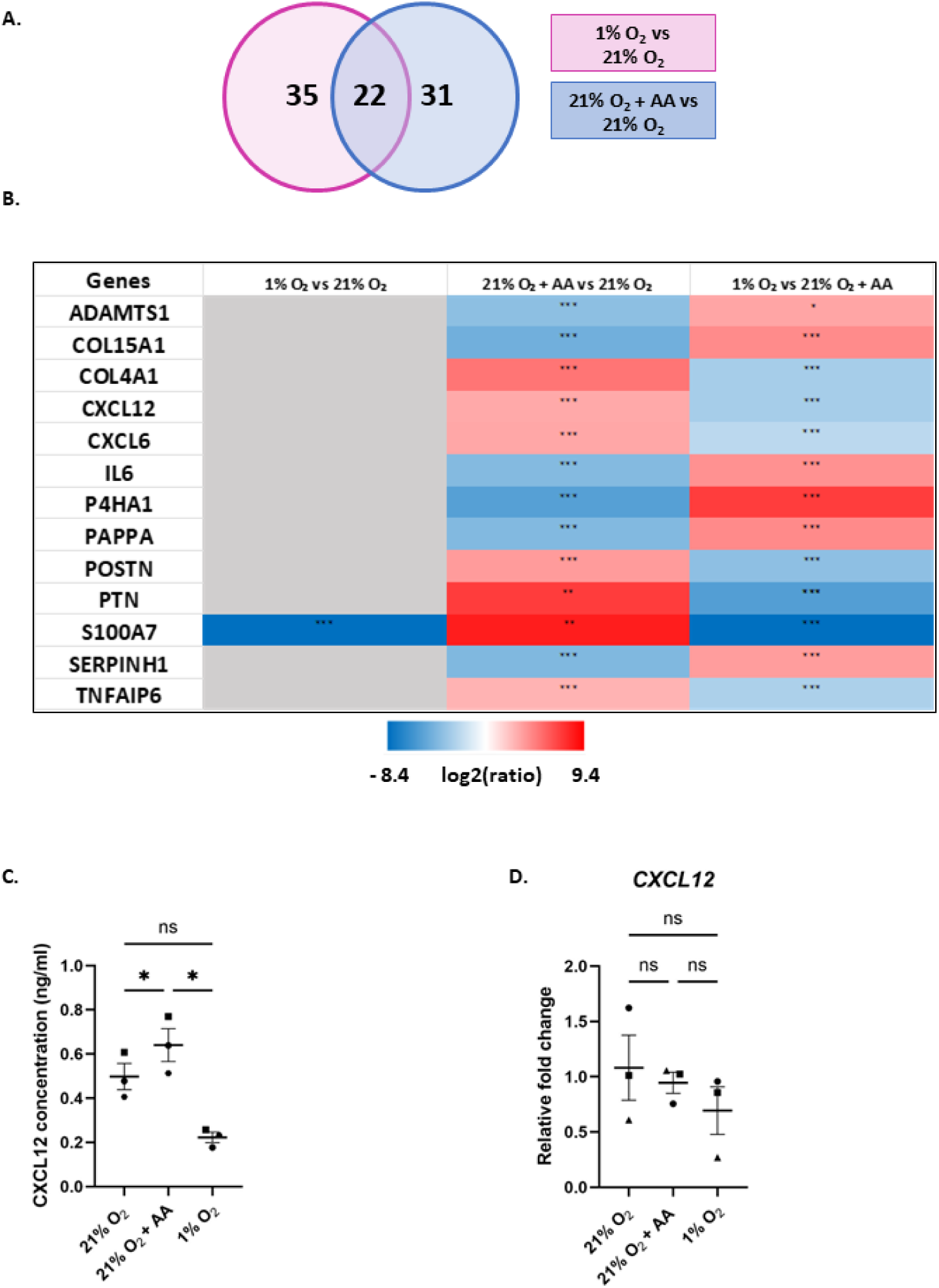
CXCL12 levels are increased in the secretome of AA-treated cells but not of hypoxic cells. Primary HDFs were cultured for 4 days in normoxia (21% O_2_) or hypoxia (1% O_2_) in the presence or absence of stabilized AA (300 µM) as indicated. **A, B.** Supernatants of HDFs were analyzed using data-independent acquisition mass spectrometry. **A.** Venn diagram showing the common and specific modifications in matrisome components when comparing the protein levels after AA treatment (21% O_2_ + AA vs 21% O_2_) or in hypoxic cell cultures (1% O_2_ vs 21% O_2_). **B.** Heat map highlighting fold changes of a selection of 13 secreted matrisome proteins between the indicated conditions. **A, B.** Protein quantification was carried out on N = 3 donors. The proteins were considered as differentially expressed between two conditions when their fold change was > 1.5 or < 0.66, with a p-value < 0.05 (* p-value < 0.05, ** p-value < 0.01, *** p-value < 0.001). **C.** ELISA test detecting CXCL12 in the supernatant of cells cultured under normoxia, with or without AA, or under hypoxia after 11 days. Statistical analysis was performed on N = 3 donors, using RM one-way ANOVA test (normal distribution of data) (ns: non-significant, * p-value < 0.05). **D.** RT-qPCR analysis of *CXCL12* transcripts in cells cultured under normoxia, with or without AA, or under hypoxia for 4 days. The results were normalized to the mean of the control condition (21% O_2_). Statistical analysis was performed on N = 3 donors, using RM one-way ANOVA test (normal distribution of data) (ns: non-significant).

### CXCL12/CXCR4 signaling is required for AA-induced collagen secretion

We next aimed to uncover the potential role of CXCL12 in AA-induced collagen I secretion. CXCL12 is known to activate the CXCR4 receptor, leading to the activation of multiple G protein-dependent downstream signaling pathways, thereby regulating diverse cellular functions such as migration, adhesion and transcriptional activation (for review [48]). In line with the reported role of CXCL12 in collagen secretion [47], we showed that the inhibition of the CXCR4 receptor by AMD3100 in AA-treated HDFs led to a marked intracellular retention of collagen I. Surprisingly however, the intracellular collagen I detected was mainly in its mature form (Figure 6A). Interestingly, this retention was fully specific to AA-induced collagen secretion as the inhibition of CXCR4 had no effect in cells cultured without AA (either normoxic or hypoxic conditions) (Figure 6B). The intracellular accumulation of collagen in AA-treated cells coupled with CXCR4 inhibition, was associated to a significant decrease in collagen deposition into the ECM (Figure 6C). Together, these results demonstrate that CXCL12/CXCR4 activation is crucial for AA-induced collagen secretion. This mechanism is specific to AA-treated cells and may explain why AA had a stronger effect on collagen secretion than hypoxia, despite its weaker induction of collagen proline 4-hydroxylation (Figures 1A and 4B). To further visualize the intracellular retention of collagen, and depict in which compartment the intracellular proteolytic maturation may occur in presence of the CXCR4 inhibitor, we imaged intracellular collagen I using super resolution microscopy. In untreated cells, collagen I was homogeneously distributed as small aggregates within the cytoplasm, likely within RER (Figure 6D). This is supported by the dilated cisternae observed by TEM (Figure 3B). Surprisingly, AA treatment induced an intracellular re-localization into aggregates accumulating next to the nucleus, possibly Golgi structures. This was accompanied by the expected formation of numerous extracellular fibrils. Strikingly, in cells treated with both AA and AMD3100, the number of extracellular fibrils was strongly reduced and, concomitantly, the collagen I labelling seemed to be localized at the plasma membrane. This suggests that collagen I is blocked near the plasma membrane when CXCL12/CXCR4 is inhibited (Figure 6D) and explains 1) the decreased extracellular deposition and 2) the intracellular accumulation of collagen observed by Western blot (Figures 6A and 6B). Together, these observations suggest that, in case of CXCR4 inhibition, collagen I is likely retained at the plasma membrane, after a transit trough the secretory pathway, where it can be proteolytically processed [49].

**Figure 6.**
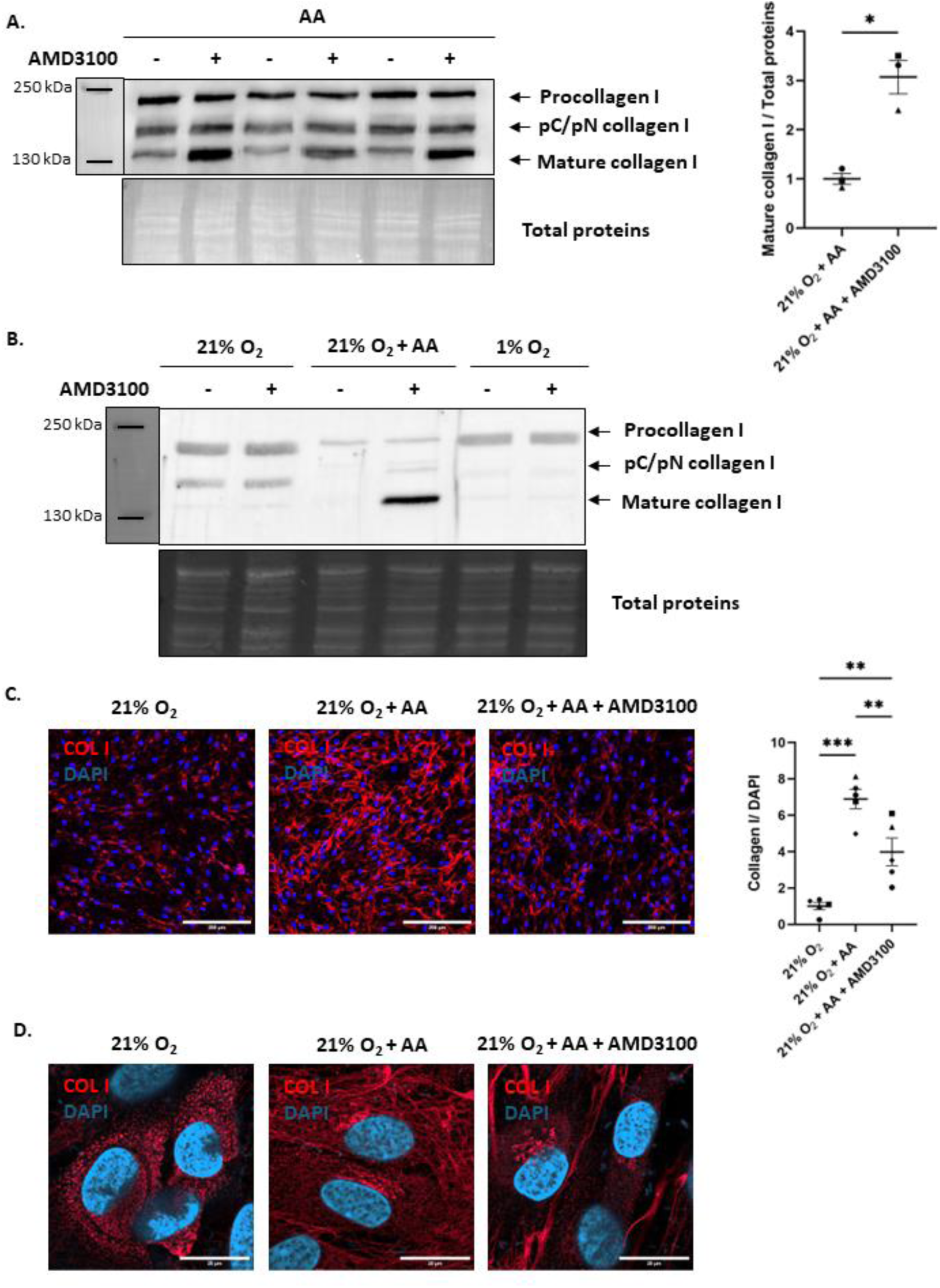
Inhibition of the CXCL12/CXCR4 axis leads to intracellular collagen retention associated with impaired collagen deposition into the matrix. **A.** Primary HDFs from three donors were cultured for two days in presence of stabilized AA (300 µM), with or without AMD3100 (10 µM). Western blot analysis of the collagen I intracellular content (reducing conditions; 6% acrylamide gel). Intracellular mature collagen I levels were normalized to total protein levels (stain-free detection) and to the mean of the control conditions (21% O_2_). Statistical analysis was performed on N = 3 donors, using paired t-test (normal distribution of data) (* p-value < 0.05). **B.** HDFs were cultured for 2 days in normoxia (21% O_2_) or hypoxia (1% O_2_) in the presence or absence of AA (300 µM) and AMD3100 (10 µM). Western blot analysis of the collagen I intracellular content (reducing conditions; 6% acrylamide gel). **C.** Immunostaining of collagen I in unpermeabilised fixed fibroblasts after 4 days of culture. Quantification was performed with ImageJ using the area covered by collagen I labelling (red) normalized to the area covered by DAPI labelling (blue) and to the mean of the control condition (21% O_2_). Statistical analysis was performed on N = 5 donors, using RM one-way ANOVA test (normal distribution of data) (** p-value < 0.01, *** p-value < 0.005). Scale bar indicate 200 µm. **D.** Immunostaining of collagen I in permeabilised fixed fibroblasts after 4 days of culture analyzed with super-resolution microscopy (Airyscan). Scale bar indicates 20 µm.

Although CXCL12/CXCR4 activation is required for AA-induced collagen I secretion, treatment of normoxic cultures with CXCL12 alone was not sufficient to alter intracellular or extracellular collagen I levels (Supplemental Figure 4A and 4B). This result is in contradiction with the observations of Patalano *et al.,* who showed that N1 cells (from a stromal nodule of benign prostatic hyperplasia) secreted higher levels of collagen I upon CXCL12 addition to the media [47]. They proposed that this effect resulted from the enhanced formation of COPII vesicles capable of carrying large cargoes, such as procollagen molecules. In their system, CXCL12 promoted the increased production of the CUL3/KLHL12 complex, thereby mediating Sec31 mono-ubiquitination, a process facilitating the formation of enlarged COPII vesicles, necessary for collagen trafficking [47]. We repeated the experiment described in this study, using MLN4924 – a NEDD8-activating Enzyme (NAE) inhibitor preventing cullin neddylation and cullin-RING ligase activation –, and confirmed that the inhibition of CULs decreased the levels of secreted collagen I (Supplemental Figure 4C). However, this was not associated to increased intracellular levels as it would be expected if the secretion was inhibited. In contrast, intracellular collagen I levels were also reduced, indicating an overall decrease in collagen production rather than a defect in collagen secretion (Supplemental Figure 4D), most likely due to off-target or toxic effects of the inhibition of all cullins. In conclusion, whilst the key role of CXCL12/CXCR4 activation in AA-induced collagen secretion is evident, the mechanism allowing the promotion of collagen I secretion remains to be elucidated.

## Discussion and conclusion

We here initially confirmed that AA and hypoxia are two independent triggers of collagen I deposition. We showed that, in addition to their common and additive ability to promote collagen hydroxylation, they also play specific roles during the subsequent steps of collagen biosynthesis, ultimately leading to different ultrastructures of collagen fibrils. This difference is mainly linked to a distinct collagen I intracellular trafficking and secretion. We uncovered a new molecular mechanism specifically engaged in the presence of AA and essential for AA-indued collagen secretion, which involves enhanced collagen intracellular trafficking, increased CXCL12 extracellular levels, and the activation of its cognate CXCR4 receptor. Indeed, CXCR4 inhibition, in the presence of AA, was associated with the retention of collagen I at the plasma membrane and impaired collagen deposition into the ECM. We below discuss our findings, focusing on the different steps of collagen biosynthesis and how they are affected by AA and hypoxia.

### Regulation of collagen transcription

The effects of the two stimuli on collagen secretion could be first explained by differences in collagen transcription. Whilst the transcription of *COL1A1* and *COL1A2* genes remained unchanged from the statistical point of view, AA increased their average transcription by ∼2 fold (Figure 2A). However, this effect is highly donor-dependent with a high increase in transcription (up to 2.6-fold for *COL1A1* and 6-fold for *COL1A2*) for some donors, but none for others. The AA-induced collagen transcription could be explained by the low intracellular levels of collagen I in presence of AA (Figures 3A and 6D). Indeed, Pinnel *et al*. suggested that the high secretion rate triggered by AA supplementation may lead to increased transcription, whilst, in absence of AA, the intracellular accumulation of collagen represses its own transcription by a negative feedback loop [50].

### Collagen hydroxylation and the extent of collagen secretion

One of the striking findings from this study is that the effect of AA on collagen secretion cannot be solely explained by the enhanced collagen proline 4-hydroxylation, as it has been commonly assumed to date. Indeed, it was discovered in the 1970s, that in absence of AA, collagen accumulates intracellularly and is strongly under-hydroxylated [51]. It was subsequently observed that AA increases both collagen proline hydroxylation and collagen secretion [52]. These observations led to the understanding that the stimulatory effect of AA on collagen secretion is mediated by its ability to promote collagen proline hydroxylation. However, our results show that hydroxylation is not the only mechanism promoting collagen secretion. Indeed, hypoxic cultures led to procollagen molecules with a higher proportion of 4-Hyp residues compared to AA-treated cells (Figure 4B), despite their weaker effect on collagen I secretion. This apparent contradiction led us to explore our secretome results in more detail and to unravel the role of CXCL12 and its cognate CXCR4 receptor in AA-induced collagen I secretion. CXCR4 inhibition not only triggered intracellular procollagen I retention, but also its premature intracellular proteolytic maturation, as observed by western-blot in Figure 6A and 6B. Super-resolution imaging of intracellular collagen distribution shed light on this finding. Upon CXCR4 inhibition, collagen I seems to accumulate, in the periphery of the cells, near the plasma membrane (Figure 6D), likely after a transit trough the secretory pathway. This is consistent with previous observations in tendon fibroblasts, showing intracellular proteolytic maturation of collagen upon disruption of its secretion [49]. Indeed, proteolytic enzymes are also known to undergo maturation in the Golgi apparatus and can therefore become active intracellularly [53]. We therefore hypothesized that, in AA-treated cells, the secretion occurs too rapidly to allow significant collagen intracellular maturation. In contrast, when the secretion is prevented, the procollagen processing enzymes (e.g. BMP-1 or ADAMTS2) may have sufficient time to act intracellularly, thereby explaining that the retained collagen is fully matured.

Even if the mechanism by which the CXCL12/CXCR4 axis drives AA-induced collagen I secretion is not fully understood, we can hypothesize that CXCR4 inhibition prevents the last steps of the extracellular release of collagen I, and, in turn, the subsequent formation of fibrils. In our system, this hypothesis is more probable than the mechanism previously proposed by Patalano *et al*., relying on the inhibition of the formation of large COPII vesicles necessary to export procollagen from the ER [47]. Indeed, the formation of large COPII vesicles is triggered by the monoubiquitinylation of Sec31 by CUL3/KLHL12 complex [54]. However, the inhibition of CUL3 with MLN4924 failed to mimic the effects of CXCR4 inhibition (Supplementary figure 4). Furthermore, we did not observe intracellular accumulation of AMD3100-induced perinuclear collagen aggregation. Instead, there was an accumulation close to the plasma membrane upon CXCR4 inhibition (Figure 6D). Finally, TEM imaging showed that the RER cisternae had a similar size in AA-supplemented cells, regardless of AMD3100 treatment (Supplementary Figure 5), making unlikely the possible RER stress due to intraluminal protein accumulation.

### AA engages CXCL12/CXCR4 signaling via a non-canonical mechanism

The role of CXCL12/CXCR4 axis in modulating collagen I secretion was specific to AA-treated cells, as no effect of the CXCR4 inhibition was observed in AA-deficient cell cultures (Figure 6B). Whilst increased levels of chemokines are usually due to an activation of their transcription, it was not the case here. This suggested that a non-canonical mechanism was activated by AA. This hypothesis was reinforced by the fact that hypoxia did not induce *CXCL12* transcription either, despite being a direct HIF target gene [55,56]. There was therefore no transcriptional activation in any culture conditions.

As CXCL12 is a crucial chemokine in many homeostatic processes, its activity is tightly controlled and regulated at different levels. In addition to transcriptional regulations, CXCL12 levels and activity can be regulated by differential mRNA splicing, posttranslational regulation through enzymatic or chemical modifications that can alter the ability of CXCL12 to bind glycosaminoglycans or its receptor. For example, NH_2_-terminal truncation can inactivate the chemokine [57]. However, a crucial mechanism regulating CXCL12 availability and activity is its interaction with other ECM components such as the glycosaminoglycans, especially heparan sulphates, found on the proteoglycans. Interestingly, based from our proteomic dataset, soluble Syndecan-4 (SDC-4) levels are low when CXCL12 levels are high (21% O_2_ + AA vs 21% O_2_: ratio=0.67 for SDC-4 after 4 days of culture). SDC-4 is primarily a transmembrane proteoglycan, which can interact with CXCL12 through its heparan sulphate groups and promote its binding to CXCR4 [58,59]. This membrane-bound SDC-4 can however undergo shedding, notably by ADAMTS-1 [60], resulting in the release of a soluble fragment into the extracellular environment [61], which is the form detected in our secretome analysis. As no transcriptional regulation of *SDC4* was observed in the different culture conditions (Supplementary figure 6), the decreased levels of soluble SDC-4 were likely due to decreased shedding. Remarkably, ADAMTS-1 levels were also low in AA-treated conditions (Figure 5B). In summary, the presence of AA is associated with low levels of ADAMTS-1 and soluble SDC-4 and possibly higher levels of membrane-bound SDC-4, which could facilitate CXCL12 binding to CXCR4 and explain the fact that CXCL12/CXCR4 activation is specific to AA-treated cells.

Overall, we established that collagen proline 4-hydroxylation is not the only mechanism by which AA promotes collagen secretion. We observed a noticeable intracellular collagen trafficking in presence of AA and we identified a key role for the CXCL12/CXCR4 activation in collagen secretion. These findings not only shed light on AA-induced mechanisms for efficient collagen deposition, but also identify new strategies for blocking collagen I secretion, and potential new avenues for antifibrotic treatments by repurposing a clinically-approved molecule.

### AMD3100 as an antifibrotic molecule?

As AMD3100 inhibits collagen I secretion in AA-treated cells, it could be considered as a potential anti-fibrotic molecule. Indeed, in the liver, *SLC23A2* (the gene encoding SVCT2, an AA transporter) expression has been shown to positively correlate with the extent of the fibrotic area, and AA was demonstrated to be necessary for collagen secretion by activated hepatic stellate cells [62]. We can therefore hypothesize that AA contributes to the excessive collagen deposition observed in fibrotic diseases, making AMD3100 a promising therapeutic strategy to control collagen secretion and deposition. Interestingly, this inhibitor has already been considered as an antifibrotic molecule, as it was shown to slow down the progression of fibrosis in several mouse models of pulmonary fibrosis [63,64]. However, in the context of renal fibrosis, AMD3100 exacerbated tubulointerstitial fibrosis in a mouse model, demonstrating a pro-fibrotic effect [65]. This pro-fibrogenic/pro-fibrotic effect of AMD3100 has also been reported in a diabetic mouse model of wound healing [66].

Overall, AMD3100 can have either anti-fibrotic or pro-fibrotic effects depending on the type of fibrosis, the organ affected, the cell type involved and the cellular microenvironment. These contradictory effects of CXCR4 inhibition highlight the complexity and heterogeneity of fibrotic diseases, which, despite sharing common features, require distinct therapeutic approaches, as the factors regulating fibrosis can differ between tissues and pathological contexts.

## Materiel and Methods

### Cell culture and treatment

All experiments were performed using primary cultures of human dermal fibroblasts (HDF) from healthy female donors (5 independent donors obtained from Biopredict or from the tissue and cell bank of the Hospices Civils de Lyon). Fibroblasts were plated at 14,000 cells/cm² in Dulbecco’s Modified Eagle’s Media (DMEM, Merck; cat# D0822 or Biowest, Nuaillé, France; cat# L0103) supplemented with 2% foetal calf serum (FCS) (Eurobio Scientific; cat# CVFSVF17-0U) and 1% antimycotic and antibiotic solution (Sigma-Aldrich; cat# A5955). Cells were treated or not with 300 µM 2-phospho-L-ascorbic acid (AA) (Sigma-Aldrich; cat# 49752) and cultured either in normoxic conditions at 21% O_2_ and 5% CO_2_ (SANYO MCO – 20AIC incubator) or in hypoxic conditions at 1% O_2_ and 5% CO_2_ (PhO2Box, Baker Ruskinn). Cells were washed three times with PBS and starved for 48h in serum-free DMEM media before the analysis of supernatants by mass spectrometry or ELISA. For inhibitor treatments, cells were incubated with 10 µM AMD3100 (pleraxifor octahydrochloride, MedChemExpress; cat# HY-50912), or 100 nM NEDD8-activating enzyme (NAE) inhibitor MLN-4924 (Sigma-Aldrich; cat# 5.05477).

### RNA extraction and gene expression analysis

Fibroblasts were plated at 14,000 cells/cm² and cultured for 4 days in 10 cm² culture plates. Cells were then harvested with trypsin (0.5 g/L) - EDTA (0.2 g/L) (Sigma-Aldrich; cat# T3924) and centrifuged for 5 min at 1000 g. The medium was discarded and the cells were resuspended in PBS before a second centrifugation step (5 min, 1000 g). The pellets were finally resuspended in RTL RNeasy Lysis Buffer (Qiagen; cat# 79216) and frozen at -20°C. RNA extraction was performed with the Qiagen RNeasy Mini Kit (Qiagen, cat# 74104) according to the manufacturer’s instructions. 500 ng of RNA were reverse-transcribed using the PrimeScript RT-PCR Kit (Takara Bio; cat# RR037A). RT-qPCR was performed using 5 ng of cDNA in a reaction volume of 10 µL using the TB GreenPremix Ex taq II Kit (Takara, cat# RR820L) and an Azure Cielo thermocycler (Azure Biosystem). The primer concentration was 0.2 µM. All qPCR reactions were performed in triplicates and *PPIA* was used as the housekeeping gene for ΔCT normalisation. The primer sequences used were as follows: *PPIA* Forward, GCTTTGGGTCCAGGAATGG, Reverse, GTTGTCCACAGTCAGCAAT; *COL1A1* Forward, GCCAAGACGAAGACATCCCA, Reverse, GGCAGTTCTTGGTCTCGTCA; *COL1A2* Forward, AAATATCGGCCCCGCTGGAA, Reverse, GGCCTTTGGGTCCAGGGAAT; *P4HA1* Forward, CAGTACATGACCCTGAGACTGGA, Reverse TGGGGTTCATACTGTCCTCCAAC; *CXCL12*Forward, AGCCTGAGCTACAGATGCCC, Reverse, AGCTTCGGGTCAATGCACAC; Reverse, TTCTTGGGTTCGGTGGGGAC. *SDC4* Forward, CGGAGCCCTACCAGACGATG,

### Western blot

Fibroblasts were plated at 14,000 cells/cm² and cultured in 60 mm² culture plates. ECM proteins were collected in Laemmli buffer (325 mM Tris-HCl pH 6.8, 25% glycerol, 10% SDS), after a decellularization step in 20 mM NH_4_OH for 40 min and several washes with PBS. Intracellular proteins were collected in RIPA buffer (Sigma-Aldrich; cat# R0278) containing a protease and phosphatase inhibitor cocktail (ThermoFisher Scientific; cat# A32961). Protein concentrations were determined using the Pierce BCA protein assay (ThermoFisher Scientific; cat# 23225) and a TECAN Spark microplate reader (Tecan). Prior to electrophoresis, 100 mM Dithiothreitol (DTT; final concentration) was added to the protein samples (intracellular proteins or supernatants). Twenty µg of proteins were loaded on 5, 6 or 10% acrylamide gels, as indicated in figure legends, and separated in running buffer (25 mM Tris-HCl pH 9, 0.2 M Glycine, 0.1% (w/v) SDS). The quality of the migration was checked with 2,2,2-trichloroethanol (0.5% (v/v) in gels; Sigma-Aldrich; cat# T54801) protein labelling after UV activation of the gels (stain-free detection). Proteins were then transferred onto ethanol-activated PVDF membranes (pore size 0.45 µm; Millipore; cat# 831.1) in 10 mM N-cyclohexyl-3-aminopropanesulfonic acid pH 11 and 10% ethanol buffer. The quality of the transfer and total proteins were assessed with stain-free detection. Membranes were blocked for 2 h, using 5% (w/v) milk and 0.05% (v/v) Tween-20 (Euromedex; cat# 2001-B) in PBS. Primary antibody incubation was performed overnight at 4°C under shaking in 5% (w/v) milk and 0.05% (v/v) Tween-20 in PBS solution. The primary antibodies used were anti-human collagen I polyclonal antibody (Novotec; cat# 20111, 1:1000 dilution), polyclonal anti-human HIF-1α antibody (Proteintech; cat# 20960-1-AP) and monoclonal anti-human HIF-2α antibody (Fortis Life Sciences; cat# A700-003). Secondary antibody incubation was performed for 1 h at RT, in 5% (w/v) milk and 0.05% (v/v) Tween-20 in PBS solution. The secondary antibody used was an anti-rabbit IgG, HRP-conjugated (Cell signaling, Danvers, MA, USA; cat# 7074, 1:10,000 dilution). Chemiluminescent detection was performed with the Western Blotting ECL™ Select kit (Cytiva; cat# RPN2235) and imaged with a Fusion FX camera (Vilber Lourmat). Quantification was performed with the ImageQuantTL software and normalized to the total protein signal obtain with stain-free detection.

### Immunofluorescent labelling

Fibroblasts were plated at 14,000 cells/cm² and cultured in Permanox 8-well Lab-Tek chamber slides (ThermoFisher Scientific; cat# 177445). Cells were washed three times in PBS, and fixed with 4% PFA in PBS. Saturation was performed in 4% BSA (Sigma-Aldrich; cat# A3059) in PBS solution for one hour at RT. Primary antibody incubation was perfomed overnight at 4°C, in a 4% (w/v) BSA in PBS solution containing the anti-collagen I rabbit polyclonal antibody (Novotec; cat# 20111, 1:200 dilution). Secondary antibody incubation was performed, for 1 h at RT, in 4% (w/v) BSA in PBS solution, containing an Alexa Fluor™ 546-conjugated anti-rabbit antibody (Invitrogen; cat# A11035; 1:1000) and DAPI (Euromedex; cat# 1050A; 1:1000). Upon 3 PBS washes, the Lab-Tek slides were mounted with Fluoroshield mounting medium (Sigma-Aldrich; cat# F6182). The images were acquired with a Nikon TiE inverted microscope, objective 20X. Image analysis was performed using ImageJ. The area covered by collagen I labelling was measured and normalized to the area covered by DAPI staining.

For super-resolution microscopy, the images were obtained using a Zeiss LSM800 Airyscan Axio Observer.Z1/7 microscope equipped with a Plan-Apochromat 63×/1.4 oil DIC objective and a GaAsP-GMT detector. Images were acquired at a resolution of 1666 × 1666 pixels, with 16-bit encoding and a pixel size of 0.035 × 0.035 μm, corresponding to a field of view of 58.81 × 58.81 μm. The different fluorophores were analyzed sequentially across three tracks. Track 1 used the 640 nm laser at 2% power, with a detection window ranging from 642 to 700 nm, a pinhole set to 5 Airy units (281 μm), and a gain of 820 V; the signal was acquired in Airyscan mode. Track 2 used the 405 nm laser at 0.6% power, with a detection window ranging from 400 to 565 nm, a pinhole set to 1 Airy unit (41 μm), and a gain of 710 V; this channel was acquired in conventional confocal mode. The acquisition time was 2.5 μs/pixel, with a line time of 9.89 ms and a frame time of 16.72 s. Scanning was unidirectional, and no averaging was applied. Images acquired in Airyscan mode were subsequently reconstructed using the automatic Airyscan 2D processing algorithm, with processing parameters of 6.6 for Track 1 and 6.9 for Track 2.

### Transmission Electronic Microscopy

Fibroblasts were plated at 14,000 cells/cm² in 6-well plates and cultured for 11 days to obtain an overconfluent cell layer. Cells were fixed with 2% glutaraldehyde (EMS) in 0.1 M sodium cacodylate (pH 7.4) buffer at RT. After 3 washes in 0.2 M sodium cacodylate buffer, cell cultures were contrasted with 0.2% (v/v) Oolong tea extract (EM grade) in the same buffer, for 1 h at RT. They were then washed and post-fixed with 1% aqueous osmium tetroxide (EMS) containing 1.5% potassium cyanoferrate for 1 h at RT. After 3 washes in 0.2 M sodium cacodylate and 1 wash in water, they were dehydrated in a graded series of ethanol at RT and embedded in Epon. After polymerization at 60°C for 72 h, ultrathin sections (100 nm) were cut on a UC7 ultramicrotome (Leica Microsystems) and collected on Nickel grids 150 mesh. Sections were stained with lead citrate for 5 min and observed with a transmission electron microscope JEOL 1400JEM (Tokyo) operating at 100kV and equipped with a camera Orius 1000 gatan and Digital Micrograph software v1.7.

### Mass Spectrometry

#### In-Solution enzymatic digestion

30 µg of protein per sample were mixed with the lysis buffer provided in the EasyPep™ Mini Kit (A40006, Thermo Scientific). The samples were reduced and alkylated by incubation at 95°C for 10 min with continuous shaking at 1000 rpm. Subsequently, the samples were digested with a Lys-C/Trypsin enzyme mixture for 3 h at 37°C, following the manufacturer’s protocol. After the cleanup step, the resulting peptides were dried, resuspended in 0.1% formic acid, and quantified using the Quantitative Fluorometric Peptide Assay (23290, Thermo Scientific).

#### NanoLC-MS/MS analysis

Samples were analyzed using a Vanquish NEO nano-LC system (Thermo Scientific) coupled online to an Orbitrap Exploris 480 mass spectrometer via an EASY-Spray ion source (Thermo Scientific). A total of 150 ng of peptide sample per condition was loaded onto a PepMap NEO C18 trap column (300 µm ID × 5 mm, 5 µm, Thermo Fisher Scientific) and subsequently separated on an Acclaim PepMap100 C18 analytical column (50 cm × 75 µm ID, 2 µm, 100 Å, Thermo Scientific). Peptides were eluted using a 60 min linear gradient from 3% to 25% of buffer B (A: 0.1% (v/v) formic acid in H₂O; B: 0.1% (v/v) formic acid in 80/20 ACN/H₂O) over 50 min, followed by an increase from 25% to 40% buffer B in 10 min, then from 40% to 100% buffer B in 1 min, held for 10 min, and finally returned to initial conditions in 10 min. The total run time was 80 min at a flow rate of 300 nL/min. The column oven temperature was maintained at 40°C.

MS data were acquired using a data-independent acquisition (DIA) strategy. Two experiments were combined. First, an MS survey scan (350–1200 Th) was performed at a resolution of 60,000 at m/z 200, with an ion target value of 3 × 10⁶ and a maximum injection time of 45 ms. Second, a DIA experiment was conducted with the following parameters: resolution of 30,000 at m/z 200, ion target value of 1 × 10⁶, and automatic maximum injection time. Twenty-seven isolation windows of 20 Th width, with a 1 Th overlap, were applied over a mass range of 350–900 Th, using a normalized HCD collision energy of 27%. The spray voltage was set to +1900 V, and the ion transfer tube temperature was maintained at 275°C.

### Data Analysis

Proteins were identified and quantified using the Spectronaut 19 software (Biognosys) in directDIA^TM^ analysis, without a library, using the Homo sapiens database (UniProt, release of July 2024; 20,518 sequences) and a contaminant database. Parameters were kept at their default values (BGS factory settings). Up to two missed cleavages were allowed. Oxidation (M, P) and acetylation (protein N-terminus) were set as variable modifications, while carbamidomethylation (C) was defined as a fixed modification. Trypsin/P was selected as the digestion enzyme parameter. Peptide and protein validations were performed at a false discovery rate (FDR) of 1%.

Protein quantification was carried out using a label-free quantitation (LFQ) approach based on fragment ion intensities. Following normalization and averaging of the abundances within each condition, protein abundance ratios were calculated. Proteins were considered differentially expressed between two conditions when their fold change exceeded 1.5 or fell below 0.66, with a p-value < 0.05.

### Proline and hydroxyproline assays

Fibroblasts were plated at 14,000 cells/cm² in 6-well plates and cultured for 4 days. Two days before harvesting the supernatants, the medium of the cells was changed for a serum-free medium. Samples consisted in 1 mL of crude serum-free culture supernatants. Samples as well as standards (prepared from a synthetic peptide containing both proline and 4-hydroxyproline residues, synthesized at JPT peptide Technologies) were dried in thermoresistant Teflon-capped microtubes and resuspended in 100 µl of 6 N HCl under anoxia in a glove box. The tubes were tightly closed and heated to 110°C overnight. One hundred µl of 6 N NaOH were added and 10 µl of the neutralized hydrolysate (or 10 µl of the 10-fold diluted neutralized hydrolysate, depending on contents) was processed for AccQTag (Waters) labelling according to the supplier’s recommendations, directly in total recovery sampler tubes. Amino-acids were analyzed by UPLC-MS on an Acquity Premier system equipped with a QDa mass detector (all from Waters), using a Cortecs C18 column (15 cm x 2.1 mm, 1.6 µm particles, Waters) and a 5-minute eluting gradient, ranging from 1% to 14% acetonitrile in water equilibrated with 0.1% (v/v) formic acid. Proline and 4-hydroxyproline were monitored based on the mass detector signal recorded in single ion monitoring in positive mode focusing at 286.2 and 302.2 Da, respectively, and at the corresponding retention times, with 10 V cone voltage. Injected sample contents were calculated in Empower using the corresponding amino-acid calibration curves based on peak areas. After data extraction, actual sample contents were corrected for sample dilutions.

### ELISA Test

Fibroblasts were plated at 14,000 cells/cm² and cultured in 12-well plates. Two days before harvesting the supernatants, the medium of the cells was changed for a serum-free medium. One mL of serum-free culture supernatants was collected after 11 days of culture, and centrifuged for 5 min at 1,000 rpm. The test was run immediately after supernatant collection using The Human CXCL12/SDF1 ELISA Kit (ABclonal; cat# USARK00266) and the experiment was conducted following the supplier’s instructions. The results were read with a TECAN Spark microplate reader.

### Statistical analysis

Statistical analyses were performed using Graph Pad Prism 10. Data normality was assessed using the Shapiro-Wilk normality test, and equality of variances was evaluated using the Brown-Forsythe test. Data were analyzed using paired statistical tests. In graphs, each fibroblast donor is represented by a different symbol. For comparison between two groups, a paired t-test was used. For the comparison involving more than two groups following a normal distribution, a repeated-measures (RM) one-way ANOVA was used when variances were equal and a RM one-way ANOVA with the Geisser-Greenhouse correction when variances were not equal. When the data distribution was not normal, groups were compared using the Friedman test, regardless of variance equality.

## Supplementary Data

**Supplementary figure 1.**
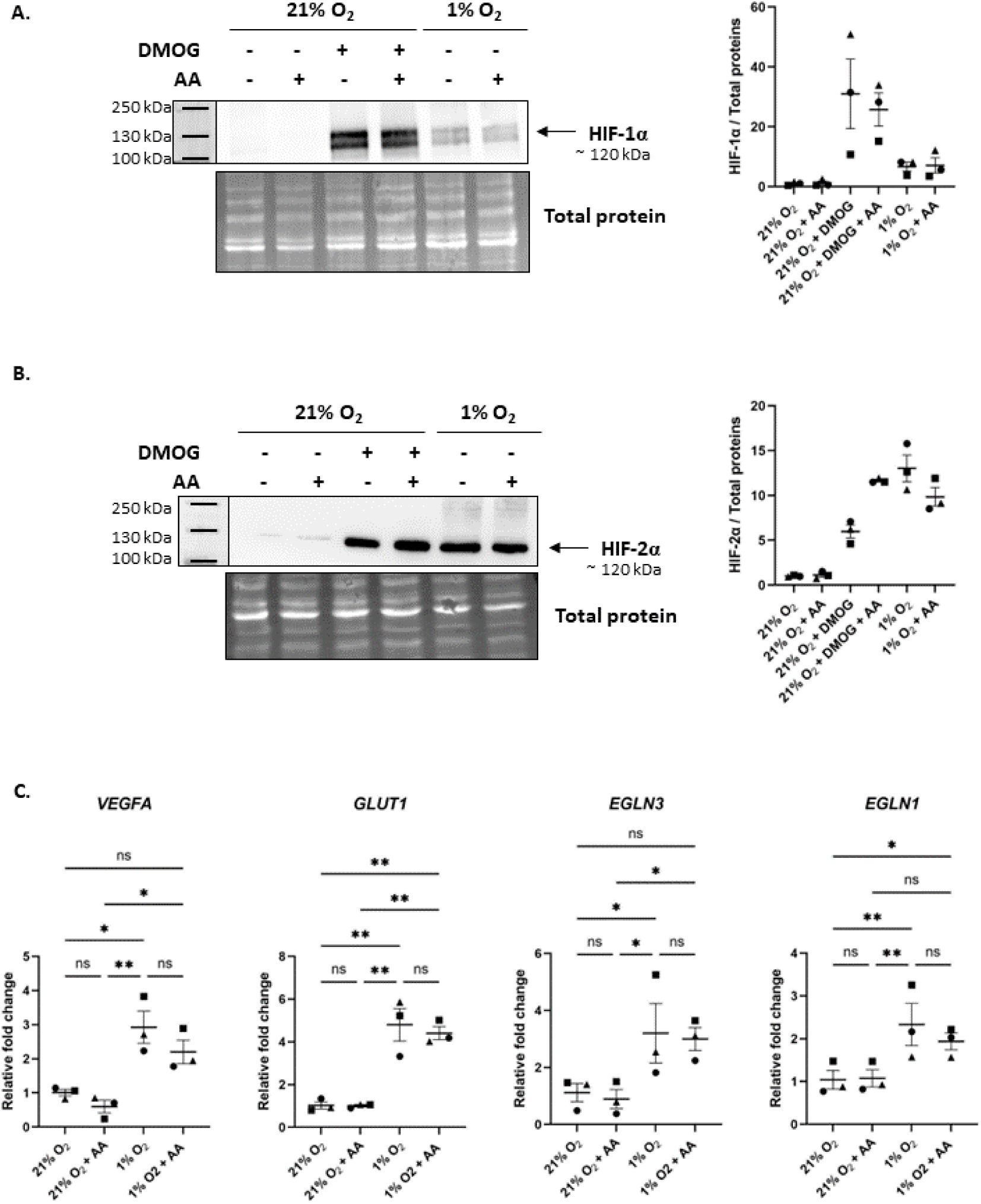
AA does not affect hypoxia-induced HIF-1α and HIF-2α stabilisation or the transcription of HIF-target genes. **A, B**. Western blot analysis of the intracellular lysates after 8 h of culture in normoxia (21% O_2_), hypoxia (1% O_2_) in presence or absence of stabilized AA (300 µM) or DMOG (0.5 µM; an HIF-P4Hs inhibitor) as indicated (reducing conditions; 10% acrylamide gel). HIF-1α and HIF-2α levels were normalized to total protein levels (stain-free detection) and to the mean of the control conditions (21% O_2_). **C.** RT-qPCR analysis of *VEGFA, GLUT1, EGLN1* and *EGLN3* transcripts after 1 day of culture. The results were normalized to the mean of the control condition (21% O_2_). Statistical analysis was performed on N = 3 donors, using RM one-way ANOVA test (normal distribution of data) (ns: non-significant, * p-value < 0.05, ** p-value < 0.01).

**Supplementary figure 2.**
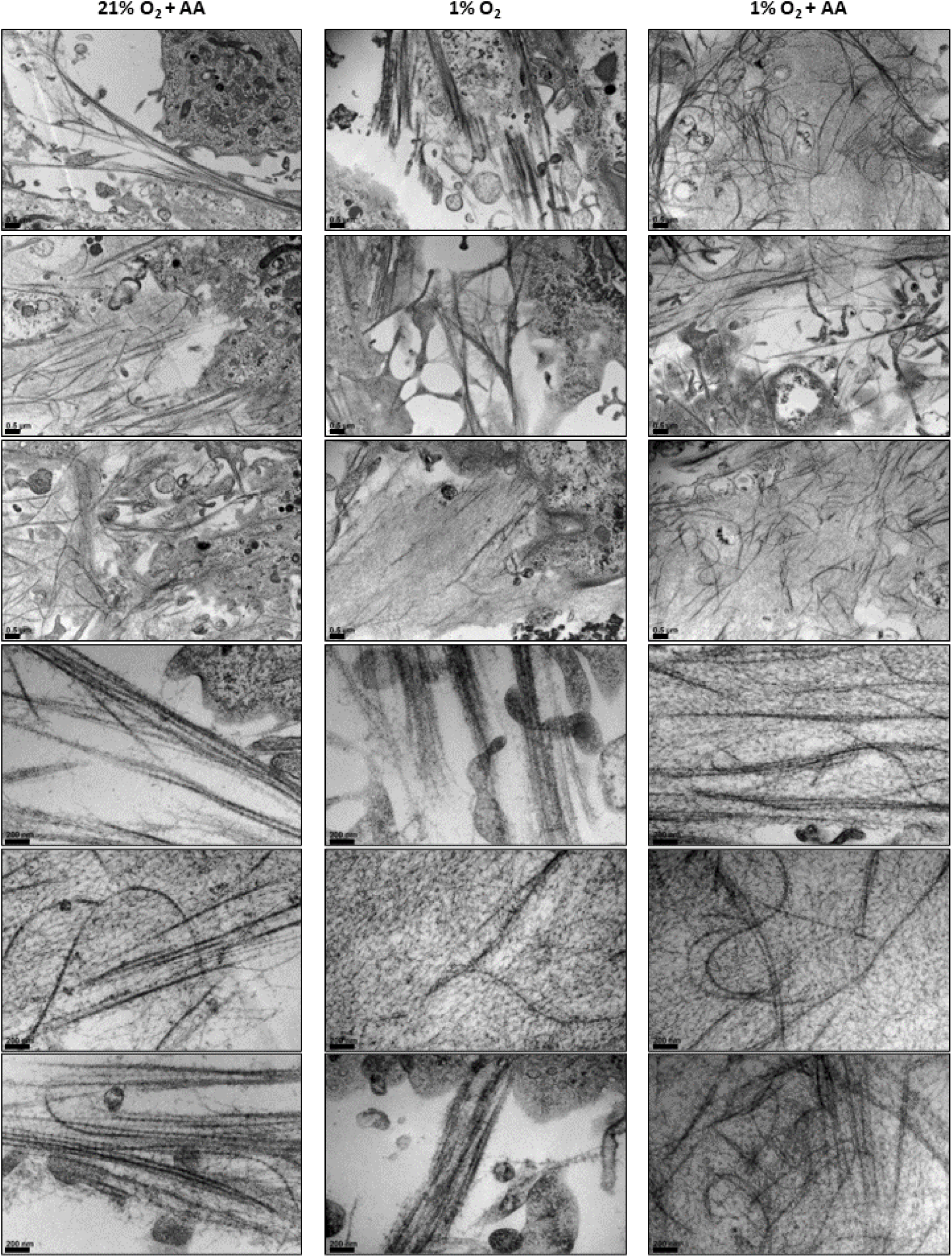
Collagen ultrastructure. ECM deposited after AA treatment or/and hypoxic exposure was imaged by TEM. Scale bar indicates 0.5 µm for the three upper rows and 200 nm for the three bottom rows.

**Supplementary figure 3.**
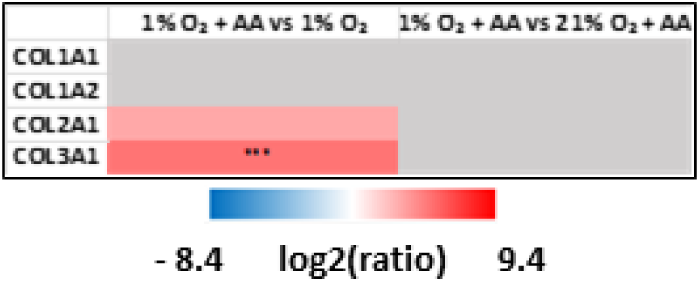
Normoxic AA-treated fibroblasts secrete similar levels of collagen I compared to hypoxic AA-treated cells. Primary HDFs were cultured in normoxia (21% O_2_), hypoxia (1% O_2_) in presence or absence of stabilized AA (300 µM) as indicated. Heat map of the mass spectrometry data for major fibrillar collagens (other conditions in Figure 2C). Protein quantification was carried out on N = 3 donors using label-free quantification. The proteins were considered as differentially expressed between two conditions when the fold change was > 1.5 or < 0.66, with a p-value < 0.05 (*** p-value < 0.001).

**Supplementary figure 4.**
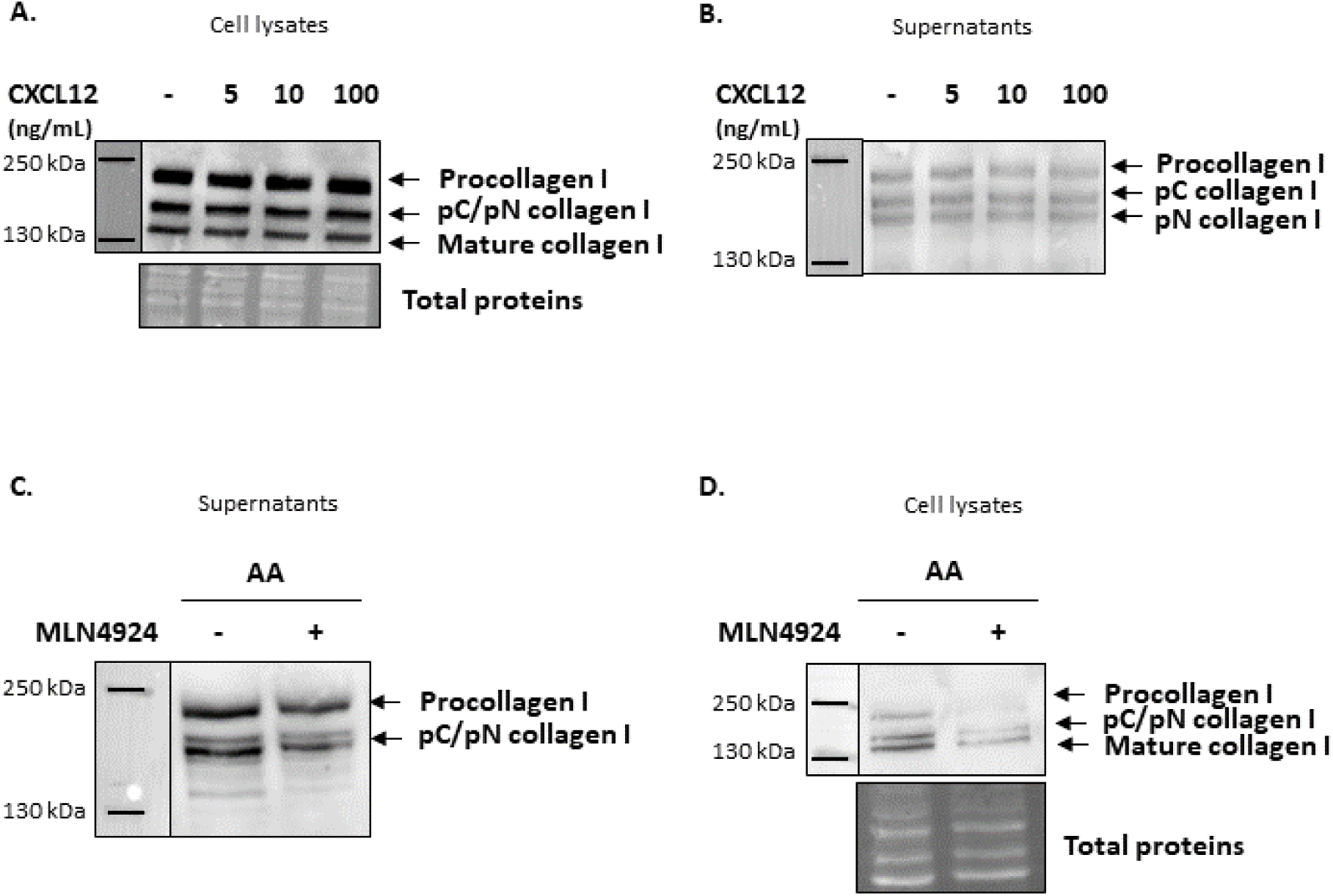
CXCL12 alone is not sufficient to promote collagen secretion. A, B. HDFs were cultured for 2 days in the presence or absence of CXCL12 (5-100 ng/mL). Western blot analysis of **A.** intracellular lysates and **B.** supernatants (reducing conditions; 6% acrylamide gel). **C, D**. HDFs were cultured for 2 days in the presence of AA and in the presence or absence of MLN4924 (100 nM) for Western blot analysis of **C.** supernatants (reducing conditions; 6% acrylamide gel) and **D**. intracellular lysates (reducing conditions; 8% acrylamide gel). Representative experiments for N = 2 donors.

**Supplementary figure 5.**
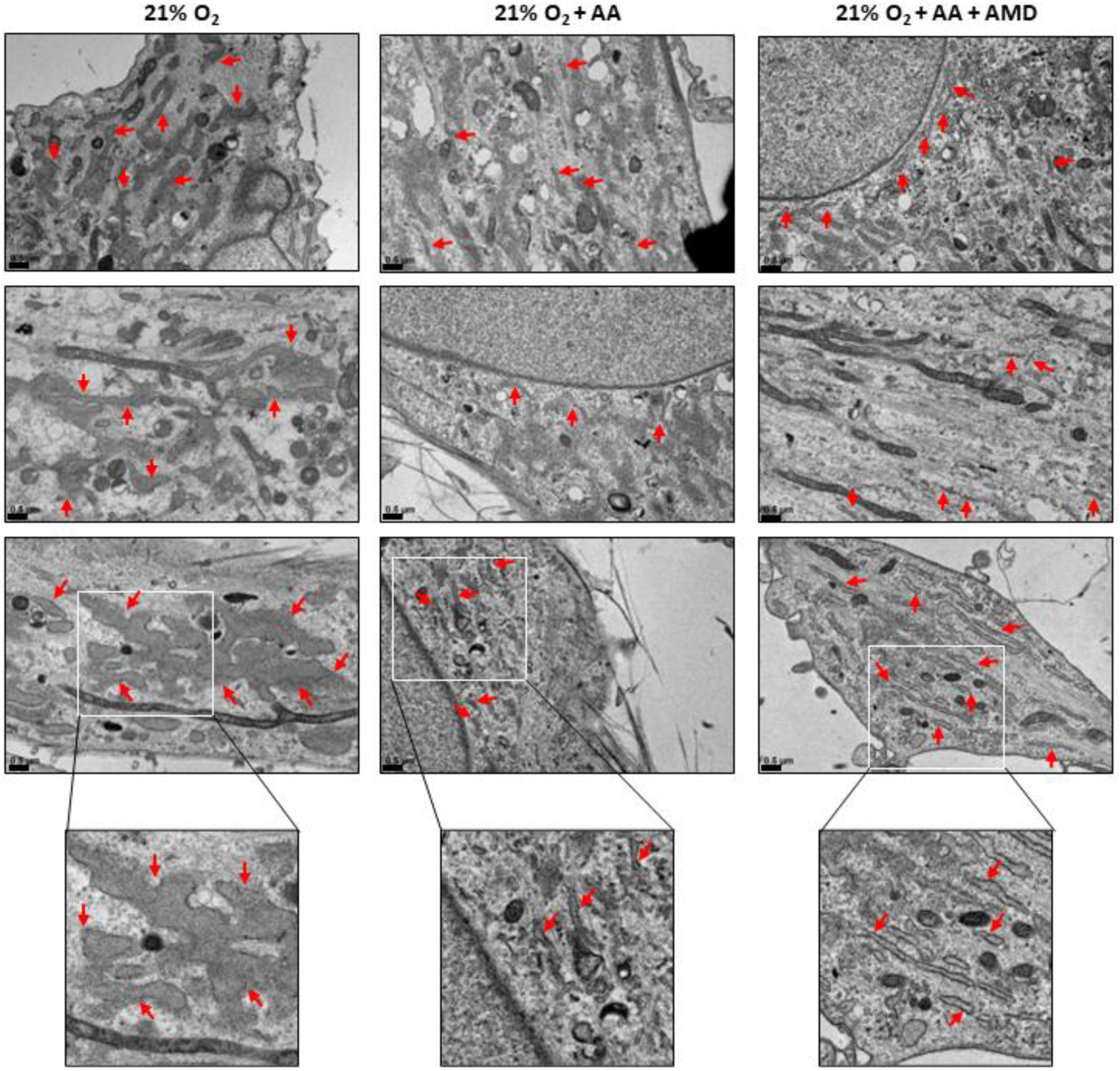
RER ultrastructure. RER shape and size after AA and AMD3100 treatments were imaged by TEM. Scale bar indicates 0.5 µm.

**Supplementary figure 6.**
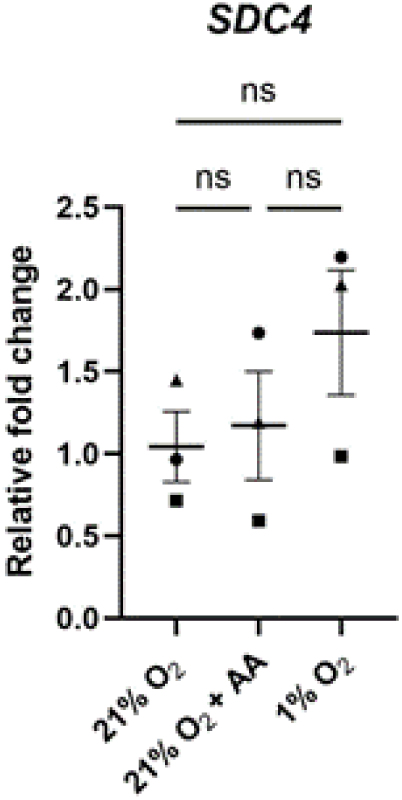
*SDC4* transcription is not affected by AA nor hypoxia. HDFs were cultured in normoxia (21% O_2_), hypoxia (1% O_2_) in the presence or absence of stabilized AA (300 µM) as indicated. RT-qPCR analysis of *SDC4* transcripts after four days of culture. The results were normalized to the mean of the control condition (21% O_2_). Statistical analysis was performed on N = 3 donors, using RM one-way ANOVA test (normal distribution of data) (ns: non-significant).

## Authors’ contributions

All cell culture, sample collection, and cell biology and biochemistry experiments were performed by MR. Proline hydroxylation measurement was performed by JBV. CB performed super-resolution microscopy. Data analysis and interpretation were done by MR, CD, SVLG, CM and VS. MR prepared the figures and drafted the manuscript under the guidance of CM and VS. VS and CM conceived and coordinated the study and wrote the manuscript. All authors reviewed drafts of the manuscript and gave final approval for publication.

## Ethics approval and consent to participate

N.A.

## Acknowledgements

MR is a recipient of the EDISS doctoral school studentship. This work was funded by Agence Nationale pour la Recherche: ANR-25-CE13–4324, from University Lyon 1 starting package (Accueil_VS) and from CNRS Innovation prematuration grant (COLCONTROL). We acknowledge the contribution of SFR Biosciences (University Lyon 1, CNRS UAR3444, Inserm US8, ENS de Lyon) Protein Science Facility, especially Frédéric Delolme and Adeline Page for the Mass spectrometry analyses. We acknowledge the support from the CNRS/IN2P3 Computing Center (Lyon - France) for providing computing and data-processing resources needed for the proteomic analysis. We thank the PrimaTiss platform (Tissue preparation and imaging platform, LBTI) and the PLATIM platform (Imaging / microscopy platform) for providing access to the microscopes, training and technical support. We also acknowledge the contribution of the CIQLE Facility from SFR Santé Lyon-Est (UAR3453 CNRS, US7 Inserm, UCBL), especially Elisabeth Errazuriz-Cerda for her help with TEM imaging and analysis.

## Abbreviations

AA: Ascorbic acid
C-P4Hs: Collagen proline 4-hydroxylases
ECM: Extracellular matrix
FCS: Fetal calf serum
ER: Endoplasmic reticulum
HDFs: Human dermal fibroblasts
HIF: Hypoxia inducible factor
NAE: NEDD8-activating enzyme
TEM: Transmission electronic microscopy
RER: Rough endoplasmic reticulum
SDC4: Syndecan-4
2-OGDD: 2-Oxoglutarate-dependent dioxygenase
4-Hyp: 4-Hydroxyproline

